# Spatial and temporal locomotor learning in mouse cerebellum

**DOI:** 10.1101/389965

**Authors:** Dana M. Darmohray, Jovin R. Jacobs, Hugo G. Marques, Megan R. Carey

## Abstract

Stable and efficient locomotion requires precise coordination of whole-body movements. Learned changes in interlimb coordination can be induced by exposure to a split-belt treadmill that imposes different speeds under each side of the body. Here we show that mice adapt to split-belt walking in a way that is remarkably similar to humans, suggesting that this form of locomotor learning is highly conserved across vertebrates. Like human learning, mouse locomotor adaptation is specific to measures of interlimb coordination, has spatial and temporal components that adapt at different rates, and is highly context-specific. Using a variety of approaches, we demonstrate that split-belt adaptation in mice specifically depends on intermediate cerebellum, but is insensitive to large lesions of cerebral cortex. Finally, cell-type specific chemogenetics combined with quantitative behavioral analysis reveal distinct neural circuit mechanisms underlying spatial *vs*. temporal components of locomotor adaptation.

## Introduction

Vertebrate control of locomotion involves multiple brain areas that work together to control its initiation, execution, and coordination (Arber, 2012; Arber and Costa, 2018; Kiehn, 2016; Orlovsky et al., 1999). For instance, basic patterns of rhythmic locomotion can be generated by spinal cord alone (Brown, 1911; Grillner and Shik, 1973; Grillner and Zangger, 1979; Sherrington, 1910), while brainstem and midbrain structures control starting and stopping, and modulate speed (Bouvier et al., 2015; Caggiano et al., 2018; Capelli et al., 2017; Esposito and Arber, 2016; Roseberry et al., 2016). Meanwhile, the cerebellum is critical for coordinating movements across the body. Uncoordinated walking, or gait ataxia, is a hallmark of cerebellar damage, and is characterized by pronounced deficits in multi-joint and interlimb coordination (Bastian et al., 1996; Hoogland et al., 2015; Machado et al., 2015; Morton, 2006; Vinueza Veloz et al., 2015)

In general, the cerebellum is thought to keep movements calibrated through a form of motor learning known as motor adaptation, an error-driven learning process through which motor responses to predictable perturbations are gradually compensated (Miles and Lisberger, 1981; Raymond and Medina, 2018; De Zeeuw and Ten Brinke, 2015). Locomotor adaptation has been demonstrated using a split-belt treadmill that imposes unequal speeds on the two sides of the body (Reisman et al., 2005; Yanagihara et al., 1993). Locomotor adaptation is cerebellum-dependent, and previous studies have shown that humans learn to adapt locomotor patterns and regain gait symmetry through specific changes in interlimb coordination (Malone et al., 2012; Morton, 2006; Yanagihara and Kondo, 1996)

Human locomotor adaptation has both spatial and temporal components – i.e., there are learned changes in both the location and timing of foot placement (Reisman et al., 2005). Many lines of behavioral evidence suggest that spatial and temporal adaptation are dissociable (Choi et al., 2009; Malone and Bastian, 2010; Malone et al., 2011; Torres-Oviedo and Bastian, 2010; Vasudevan et al., 2011). Space and time are adapted at different rates, have different developmental onsets, and are differentially influenced by explicit strategies and distraction (Malone and Bastian, 2010; Malone et al., 2012; Vasudevan et al., 2011).

Despite the abundance of evidence for dissociability of spatial and temporal locomotor learning, little is known about how this is achieved on a circuit level. Insights into specific circuit mechanisms for motor adaptation and cerebellum-dependent learning have mainly come from investigations of relatively simple behaviors like eye movements and classical eyeblink conditioning (Carey, 2011; Halverson et al., 2015; Kalmbach et al., 2011; Ke et al., 2009; Medina et al., 2000, 2002; Nguyen-Vu et al., 2013; Ohmae and Medina, 2015; Raymond et al., 1996; De Zeeuw and Yeo, 2005), due in large part to the difficulties associated with measuring whole-body movements like locomotion in animal models (Berman, 2018; Brooks and Dunnett, 2009; Brown and De Bivort, 2018; Wiltschko et al., 2015).

Here, we developed a transparent split-belt treadmill for mice that provides high-resolution, quantitative readouts of locomotor coordination (Machado, Darmohray et al., 2015), in order to study the neural circuit mechanisms underlying locomotor adaptation. We show that mouse locomotor adaptation is remarkably similar to that of humans. Further, split-belt adaptation in mice depends on an intact cerebellum, but is independent of cerebral cortex. Using a cell-type specific chemogenetic strategy, we identify for the first time a specific subregion of the cerebellum that is required for split-belt treadmill adaptation and reveal distinct circuit mechanisms underlying spatial *vs*. temporal components of locomotor learning.

## Results

### Split-belt locomotor adaptation in mice

We used a custom-built, fully transparent split-belt treadmill to assess locomotor learning in mice. The setup was a modified version of the LocoMouse setup we previously described (Machado, Darmohray et al., 2015). Animals were placed in a 4 x 30 cm acrylic corridor and two independent motors controlled the speed of transparent treadmill belts underneath each side of the body (Fig 1A). A 45-degree angled mirror below the corridor allowed both side and bottom views to be captured using a single high-speed camera (AVT Pike F-032B, 636 x 279, 330fps). Body features (paws, nose and tail) were tracked using the *LocoMouse* tracking algorithm (Fig 1B; Machado, Darmohray et al., 2015; https://github.com/careylab/LocoMouse);

**Figure 1.**
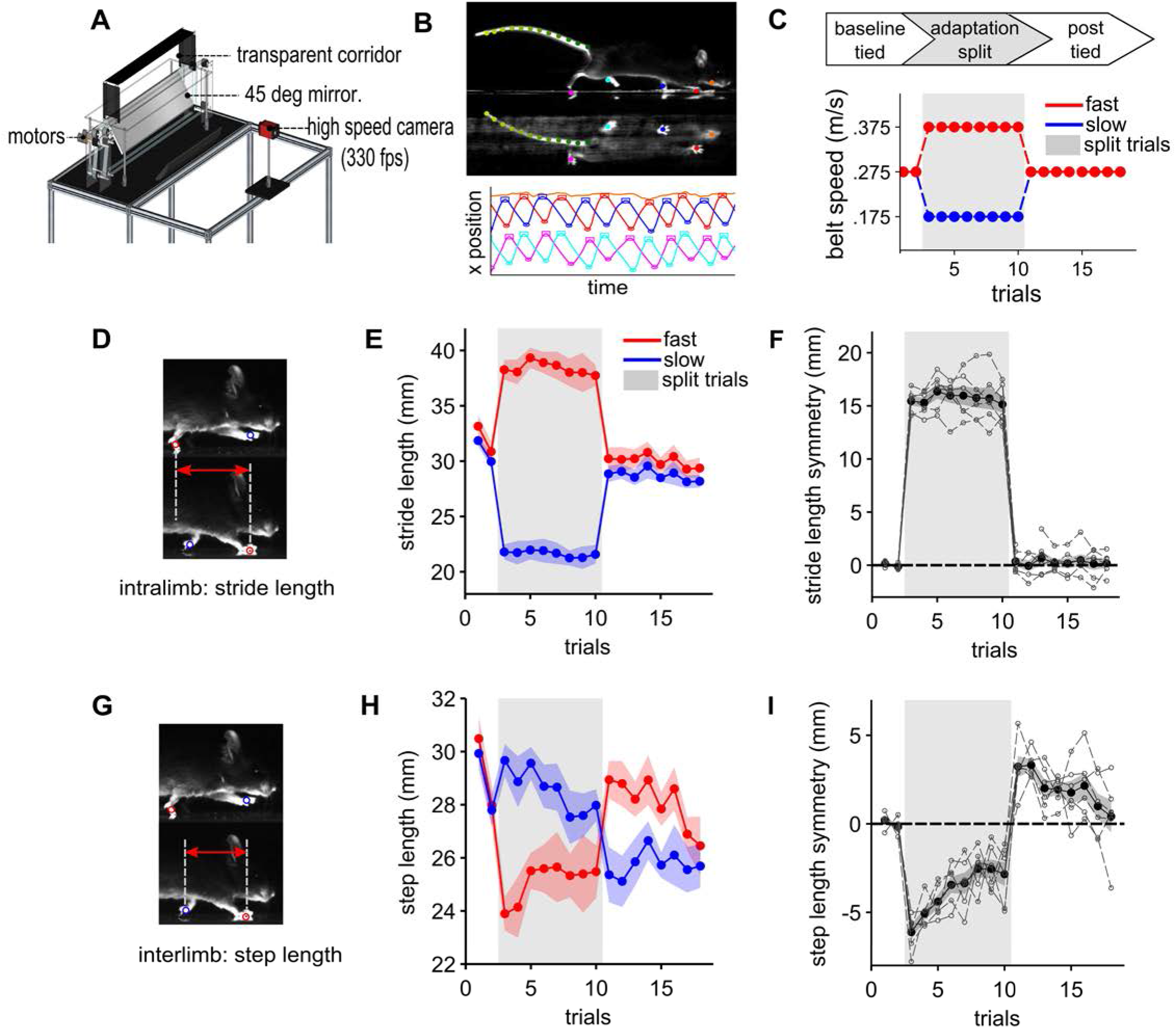
Locomotor learning on a split-belt treadmill in mice. A) Schematic of a transparent, split-belt treadmill for mice. Mice freely walk in the transparent corridor where two motor-driven belts independently control the speed of limbs on opposite sides of the body. A 45-deg. angled mirror below allows for the simultaneous capture of bottom and side views using a single high-speed camera. B) LocoMouse machine learning algorithm tracks paws, nose and tail in 3D (top). Example traces of forward (x) position versus time for the four paws and nose (bottom). C) Experimental protocol for mouse split-belt locomotor adaptation with one-minute trials. Each experiment starts with baseline trials, in which the belts are ‘tied’ (both moving at the same speed). Belts are then abruptly split at a 2.14:1 speed ratio for the adaptation phase (gray shaded area), followed by a ‘washout’ period of tied belt trials. D) The individual-limb parameter ‘stride length’ is the displacement of an individual limb from lift-off to stance onset. E) Average stride length for fast (red) and slow (blue) front limbs over a single-session split-belt experiment. F) Average front limb stride length symmetry over split-belt experiment. Individual animal measures (n = 8) are represented with open symbols and dotted lines. Group average +/-SEM are indicated by closed symbols + shadow. G) The interlimb parameter ‘step length’ for each limb is the anterior-posterior displacement between it and the contralateral limb at its stance onset. H) Average step length for fast and slow front limbs over a split-belt experiment, plotted as in E. I) Step length symmetry gradually recovers during the split-belt period and exhibits oppositely-directed aftereffects during the washout period (plotted as in F).

Freely-moving adult wild-type mice underwent adaptation protocols consisting of baseline, split-belt (adaptation), and washout phases. Sessions typically consisted of 10-23x 1-minute trials, run in rapid succession with brief rest periods (in which the motors were off) between trials. Belt speeds were equal (‘tied’) in baseline trials and were then abruptly split (approximately 2:1 speed ratio) for the adaptation phase, before returning to the original, symmetrical tied-belt speed in the washout phase (Fig 1C). Mice were habituated to the setup and treadmill walking prior to split-belt adaptation and ran naturally and without reinforcement during the experiment (Supp. Movie 1).

Previous studies have demonstrated that human split-belt locomotor adaptation is specific to measures of inter-limb coordination (Reisman et al., 2005). While individual limb parameters reset abruptly upon changes in belt speeds, interlimb measures exhibit gradual changes to regain symmetry between the legs during the split-belt period and show subsequent after-effects that gradually disappear or ‘wash out’ during the washout phase.

We first measured stride length, an individual limb parameter that measures the total excursion of the limb during a stride cycle (period from stance to subsequent stance onset; Fig 1D). Stride length scales with walking speed and as expected, mice rapidly adjusted their stride lengths to compensate for the imposed speeds under each limb, with slow and fast limbs taking shorter and longer strides, respectively (Fig. 1E). Like humans (Reisman et al., 2005), mice scaled their stride lengths with belt speed (*F_(3,5)_* = 457.34, *p* < .001), but didn’t adapt stride length symmetry over the split-belt trials (change over split, front limbs: *t*_(5)_ = 0.46, *p* = .66; Fig. 1F). Nor did they show significant aftereffects in the washout period (*t*_(5)_ = −.83, *p* = .44; Fig. 1F).

We next looked for evidence of split-belt locomotor adaptation by quantifying the interlimb parameter step length, or anterior-posterior displacement between left and right limbs at stance onset for each limb (Fig. 1G). Front limb step lengths were nearly symmetrical at baseline but became asymmetric at the start of the split-belt phase, with slow limbs taking larger step lengths than fast limbs (Fig. 1H). Step length symmetry adapted over the course of split-belt trials with mice recovering an average of 57.9% of their initial error over 8 minutes (Fig. 1I; repeated measures ANOVA for experimental phase: *F_(3,5)_* = 262.87, *p* < .001; change over split: *t*_(5)_ = −4.07, *p* = .01). Prominent after-effects were observed when belts were returned to equal speeds in washout trials (Fig 1I; *t*_(5)_ = −5.37, *p* = .003). Similar patterns of individual and interlimb changes were observed regardless of which side of the body experienced the faster belt speed, suggesting no effect of handedness (Supp. Fig. 1A; stride length scaling: *t*_(9)_ = 1.74, *p* = .12; change over split: *t*_(9)_ = .087, *p* = .93, after-effect: *t*_(9)_ = .035, *p* = .97)

Interestingly, while front limbs exhibited clear step length adaptation, evidence for learned changes in hindlimb step length was weaker. Over the split-belt period, only 19.92% of the initial asymmetry was recovered, and hind limb after-effects did not reach statistical significance (Supp. Fig. 1B, main effect of experimental phase *F_(3, 5)_* = 118.4, *p* < .001; change over split: *t*_(5)_ = −2.8, *p* = .04; aftereffect: *t*_(5)_ = −1.6, *p* = .17). Split-belt walking also affected nose and tail movements. During normal forward locomotion, mice keep their nose and tail centered nearly perfectly (Machado, Darmohray et al., 2015). In the split-belt phase, there was a tendency toward more leftward nose position (toward slow belt), with a small tendency to a more rightward position during early washout. Despite this trend, there was no significant difference in these small nose position changes across the experimental phases (Supp. Fig. 1C, *F_(4, 3)_* = 2.13, *p* = .14). Tail movements followed a similar pattern (Supp. Fig. 1D); the only significant changes across experimental phase were between baseline and early split (for middle tail segment, across experiment: *F_(4, 3)_* = 16.16, *p* < .0002; baseline to early split: *t*_(4)_ = 3.4, *p* = .027).

### Mouse split-belt adaptation does not generalize across sensorimotor contexts and does not exhibit savings

To more fully characterize mouse split-belt treadmill adaptation, we used a longer protocol designed to elicit more complete adaptation (Supp. Fig. 2A). To avoid fatigue, we divided this longer adaptation into multiple sessions. After individual sessions consisting of 7-10 minutes of split-belt walking with no washout phase, mice were returned to their home cages, and the adaptation protocol was continued on the next day. This also allowed us to investigate overnight retention of learning. Analysis of session-to-session changes revealed high retention (step length session start - previous session end: *t*_(16)_= 1.14, *p* = .27). Further, after 30 mins of total adaptation time over five days, there was near complete (83.9%) front limb step length adaptation (*F_(3, 16)_* = 98.25, *p* < .001; change from early to late split, *t*_(16)_= −5.52, *p* < .001), and prominent after-effects in tied belt trials during washout (Supp. Fig. 2A; *t*_(16)_= −13.04, *p* < .001).

Next, we addressed the context-specificity of mouse locomotor learning by examining the extent to which after-effects transferred to another locomotor context, overground walking. Instead of undergoing washout, animals were moved to a different walking context immediately following the adaptation phase (Supp. Fig. 2B). We compared step length symmetry values in the two contexts during baseline trials and after adaptation. Mice showed prominent after-effects relative to baseline on the treadmill (*t*_(4)_ = −3.39, *p* = .03) but not in overground walking (Supp. Fig 2C; OG baseline v OG adapted (*t*_(4)_ = −0.437, *p* = .66). Thus, split-belt locomotor learning in mice, as in humans, does not generalize well across sensorimotor contexts (Malone, 2011).

**Figure 2.**
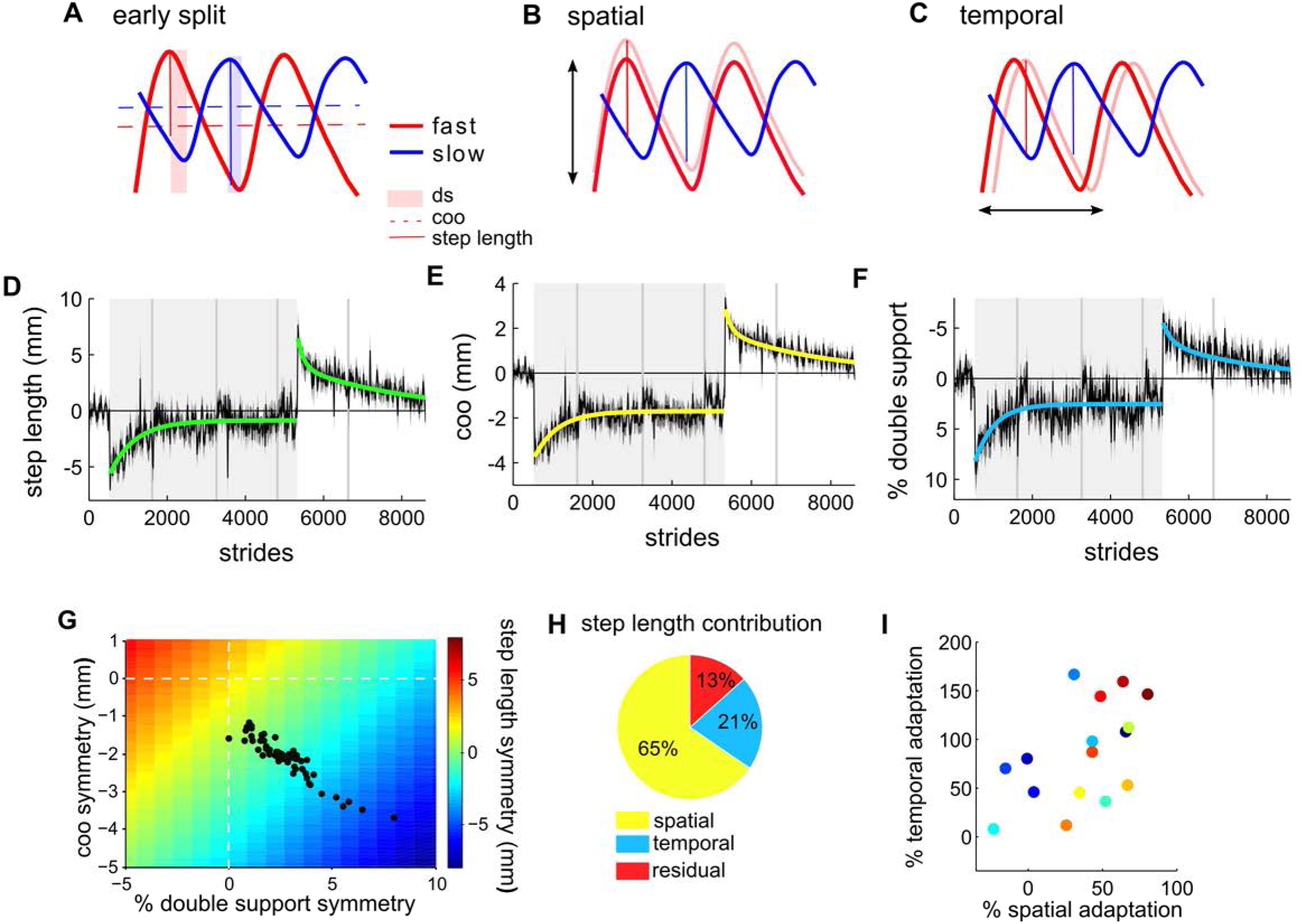
Step length adaptation has spatial and temporal components. A) Schematic of front limb stride cycles for early split-belt period, demonstrating spatial and temporal asymmetries for fast (red) and slow (blue) limbs. In early split, step length shows marked asymmetries between the limbs (vertical solid lines). This step length asymmetry is due to both spatial (center of oscillation (coo), horizontal dashed lines) and temporal (double support (ds), shaded vertical patches) asymmetries. Animals can adapt step length either spatially, by shifting the center of oscillation of the limbs (B) or temporally (C), by changing the relative timing between the limbs. (D) Average (n = 17) front limb step length symmetry over a 30-minute total split-belt period, with exponential decay fits for adaptation and washout periods. Gray shaded patch indicates split-belt trials; darker vertical lines show session breaks. Error bars represent SEM. E) Same as D for spatial parameter, center of oscillation. (F) Same as D for temporal parameter, double support. G) Simulated step length symmetry mapping based on different combinations of spatiotemporal shifts. Color code represents step length symmetry (green is symmetrical). Dashed white lines represent perfect temporal (x) and spatial (y) symmetry. Black dots represent the average step length symmetry for individual trials over the adaptation phase (data from D). In this configuration of belt speeds, our model estimates a 2 mm and 0.24 mm contribution to step length symmetry for every 1 unit change in coo (mm) or double support (% stride cycle), respectively. H) Average contribution of spatial and temporal adaptation to step length symmetry, based on the model fit to the data. I) Percent spatial and temporal adaptation for individual animals.

Many forms of sensorimotor adaptation show evidence of savings, or faster relearning upon repeated exposures to an experimental perturbation (Brashers-Krug et al., 1996; Krakauer, 2005; Medina et al., 2001). Savings is also observed in human split-belt locomotor adaptation (Malone et al., 2011). To test for savings in mouse locomotor adaptation, mice (n = 9) underwent a second round of split-belt treadmill adaptation following the near complete adaptation and deadaptation elicited by the longer, 30-min adaptation protocol. We found no difference between adaptation and re-adaptation across any phase of the experiment (Supp. Fig. 2D-F, *F_(2, 8)_* = .65, *p* = .53), suggesting no evidence for savings in mice in this task.

### Step length adaptation has spatial and temporal components

Step length is a global measure of spatiotemporal gait changes that occur during split-belt walking (Malone et al., 2012; Reisman et al., 2005). In early split-belt trials, asymmetries in both space and time contribute to overall step length asymmetry (Fig. 2A). Spatial asymmetry in step length reflects fast limb oscillation about an axis (termed center of oscillation) further back on the body than the slow limb (red and blue dashed horizontal lines, Fig 2A). Temporal asymmetries are also present in early split, with fast limbs swinging earlier relative to touchdown of the slow limb than in baseline tied walking. This temporal asymmetry results in asymmetric patterns of double support (phases of the stride cycle in which both homologous limbs are in stance; red and blue vertical shaded patches, Fig. 2A). Since individual limb parameters (eg., stride length) don’t adapt, the initial step length asymmetry (Fig 2A vertical solid lines) can be equalized either by incrementally adjusting the spatial excursions of the limbs relative to the body (spatial adaptation, Fig 2B) or by adjusting the relative timing between limb movements (temporal adaptation, Fig 2C). In humans, step length adaptation has been shown to have both spatial and temporal components (Reisman, 2005).

We quantified spatial adaptation using the symmetry of center of oscillation (COO), or the midpoint of swing and stance positions relative to the body (Fig. 2A, dashed horizontal lines). Temporal adaptation was quantified as changes in double support symmetry: the percentage of the stride duration in which two homologous limbs overlap in stance (Fig. 2A, shaded boxes). For these analyses, we analyzed step length and its spatial and temporal components over shorter (6 s) bins in the long adaptation experiment (Fig. 2D-F). Mice adapted both spatial (Fig. 2E; *F_(3, 16)_* = 145.71, *p* < .001; change over split: *t*_(16)_= −5.64, *p* < .001), and temporal (Fig. 2F; *F_(3, 16)_* = 65.56, *p* < .001; change over split: *t*_(16)_= 4.80, *p* < .001) measures during the split-belt phase and showed negative after-effects in washout tied-belt trials (Fig. 2 E,F; spatial: *t*_(16)_ = −11.59, *p* = <.0001; temporal: *t*_(16)_ = 8.18, *p* < .0001). Average adaptation and washout curves for each parameter were then fit with exponential decay functions to compare adaptation and washout rates for spatial and temporal parameters. As in human locomotor adaptation (Malone et al., 2011), exponential fits showed differential adaptation rates for spatial and temporal adaptation, with temporal adaptation occurring faster (λ = 494.4 strides) than spatial (λ = 590.0 strides). Due to rapid initial de-adaptation, the washout period was better fit with double exponential equations (Smith et al., 2006), consistent with the finding of faster washout than acquisition in human split-belt walking (Malone and Bastian, 2010).

To better understand the relationship of spatial and temporal contributions to step length adaptation, we simulated strides with triangle waveforms that were matched to average individual limb measures for slow and fast limbs (see Methods). We then shifted the waveforms for each front limb in space and time relative to each other and calculated step length symmetry for each shift. This generated a full mapping of step length error space during early split-belt walking (Fig. 2G). The simulated spatio-temporal shifts accurately predicted step length adaptation curves (r^2^ =.83).

The space of spatiotemporal combinations (Fig. 2G) shows many ways to reach step length symmetry (Fig. 2G, green). We found that animals did so in a way that simultaneously equalized all three parameters (Fig. 2G, black dots). Center of oscillation and double support are in different units, but our simulated data suggests that temporal shifts have a smaller impact on step length symmetry than spatial changes (Fig. 2H; Finley et al., 2015). Finally, individual animals show variability in their amount of overall adaptation but don’t seem to categorize into spatial or temporal groups; the best adapters adapt more of both spatial and temporal parameters (Fig. 2I).

### Locomotor adaptation requires cerebellum but not sensorimotor cortex

Previous work in humans and cats has implicated the cerebellum in split-belt treadmill adaptation (Morton, 2006; Yanagihara and Kondo, 1996). We took a number of approaches to investigate the necessity and sufficiency for the cerebellum in mouse locomotor adaptation.

First, we tested locomotor adaptation in two well-characterized ataxic mouse lines with abnormal cerebella, *reelers* and *Purkinje cell degeneration (pcd*) (Lalonde and Strazielle, 2007). *Reeler* mice have many brain abnormalities and exhibit severe ataxia that has been attributed to cerebellar maldevelopment, including cerebellar hypoplasia and lack of cerebellar folds (Fig 3A). *Pcd* is a recessive mutant mouse with moderate ataxia resulting from complete postnatal loss of Purkinje cells and subsequent partial loss of cerebellar granule cells (Fig 3E). We ran mutants and their littermate controls in a split-belt protocol using lower overall speeds than those used in the previous experiments but maintaining a similar belt ratio.

**Figure 3.**
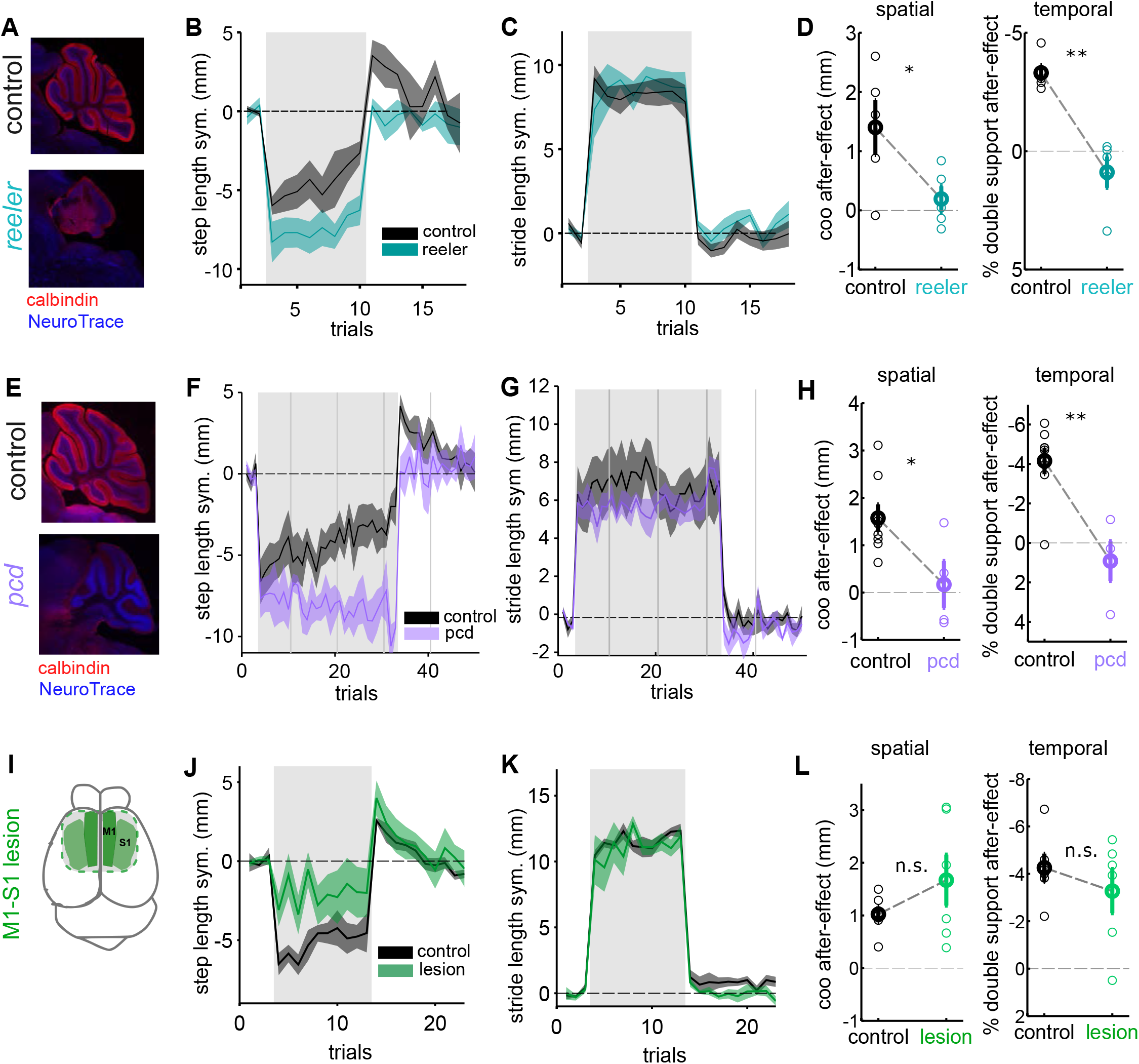
Mouse locomotor adaptation requires the cerebellum but not somatomotor cortex. A) Cerebellar sagittal section from littermate control (*top*) and reeler mutant (*bottom*) with calbindin (red) and Neurotrace staining (blue). B) Average step length adaptation curves (± s.e.m.) for reeler mutants (blue, n = 5) and littermate controls (black, n = 5). C) Mean stride length for reeler mutants and littermate controls. D) Mean after-effect (first washout trial) for center of oscillation (spatial, left) and % double support (temporal, right) for reelers and littermate controls. Each circle represents an individual animal. Thick circles are group averages (± s.e.m.). E) Cerebellar sagittal section for control and Purkinje cell degeneration (*pcd*) mice with calbindin (red, Purkinje cells) and Neurotrace staining (blue). F) Average step length adaptation curves (± s.e.m.) for *pcd* mice (purple, n = 4) and their littermate controls (black, n = 8), over 30 minutes of split-belt walking. Light gray shaded patch represents split-belt trials; gray vertical lines indicate session breaks. G) Mean stride length symmetry for pcd and littermate controls. H) Average spatial and temporal after-effects for *pcd* and littermate controls, plotted as in F. I) Schematic of aspiration zones for somatomotor cortex lesions. J) Mean step length adaptation curves (± s.e.m.) for pre-lesion control (black) and somatomotor cortex lesioned mice (green, n = 5). K) Mean stride length (± s.e.m.) for control and somatomotor lesion mice. L) Average spatial and temporal after-effects for pre- and post-lesion mice, plotted as in F. In L, significance was assessed with paired *t*-tests, otherwise, unpaired.

Despite their ataxia, reeler mice were able to respond similarly to controls to the abrupt change in belt speeds at the onset of split-belt trials (Fig 3C, first split, *reeler* vs. littermates: *t*_(4)_ = 0.82, I = .45). However, reeler mutants were impaired at locomotor adaptation compared to their littermate controls (Fig 3B, step length after-effect, *reeler* vs. littermates: *t*_(4)_ = 4.75, *p* = 0.009). Both spatial and temporal adaptation were eliminated (Fig 3D; coo aftereffect:_(4)_ = 3.501, *p* = 0.025; double support aftereffect, *reeler* vs. littermates: *t* (4) = 6.20, *p* = 0.003).

Similarly, while *pcd* mice showed normal asymmetries in both stride (Fig 3G, first split, *pcd* vs. littermates: *t*_(10)_ = .30, *p* = .77) and step length upon initial exposure to the split-belt condition, they were unable to adapt, even over three full sessions of split-belt walking (Fig 3F; after-effect compared to 0 mean distribution: *t*_(3)_ = 0.14, *p* = 0.89; aftereffect compared to littermate controls: *t*_(10)_ = −4.18, *p* = 0.002). Again, both spatial and temporal components of learning were eliminated (Fig 3H; after-effects compared to littermates: spatial: *t*_(10)_ = −2.62, *p* < 0.02; temporal: *t*_(10)_ = 4.28, *p* < 0.002).

Next, we investigated a possible role for sensorimotor cortex in split-belt treadmill (Mathis et al., 2017). Mice underwent split-belt treadmill adaptation before and after bilateral aspiration of forelimb primary motor and somatosensory cortices (Fig 3I; Supp. Fig 3). Lesioned animals, like the cerebellar mutants, were able to walk normally in the tied-belt condition and adjust their stride lengths with the change in belt speeds (Fig. 3K). Despite smaller initial split-belt asymmetries in these animals, however, the normal aftereffects indicated that locomotor adaptation itself was unaffected by the lesion (pre-post lesion step length aftereffect: Fig 3J; *t*_(5)_ = −1.17, *p* = 0.29). Both spatial and temporal components of learning were intact (Fig 3L; spatial: *t*_(5)_ = −1.13, *p* = .31; temporal: *t*_(5)_ =-1.01, *p* = 0.36).

### A cerebellar region for locomotor adaptation

The results of Fig. 3 are consistent with findings from human patients in implicating a critical role for the cerebellum in split-belt treadmill adaptation (Morton & Bastian, 2006). However, it is not known how different cerebellar regions contribute to learning. Localizing locomotor adaptation to a particular region or regions of cerebellar cortex is complicated by its highly convoluted surface and fractured somatotopy (Manni and Petrosini, 2004). In contrast, all output from the cerebellar cortex goes through Purkinje cells that project to one of a set of deep cerebellar (and/or vestibular) nuclei. Within the context of locomotion, each of the cerebellar nuclei have the potential to impact spinal locomotor circuitry, through direct or indirect projections (Chambers and Sprague, 1955; Morton and Bastian, 2007). To begin to narrow down the potential sites of cerebellar plasticity for split-belt locomotor adaptation, we targeted each deep cerebellar nucleus (medial, interposed and lateral) using a chemogenetic approach (Armbruster et al., 2007; Roth, 2017).

We chose a chemogenetic approach because the time scale of DREADD action is well aligned with that of our task, in which learning evolves over a time course of minutes, with no particular trial structure or corresponding discrete events (Roth, 2016, 2017; Wolff and Ölveczky, 2018). Further, this method, in contrast to more acute manipulations like optogenetics, did not impair baseline walking or cause overt ataxia that could confound motor learning and motor coordination (Supp. Movie 2).

We injected retrograde adeno-associated virus (Tervo et al., 2016) unilaterally into the cerebellar nuclei of L7-cre mice for the cre-dependent expression of the Gi-coupled (inhibitory) DREADD, hM4Di, in Purkinje cells terminating there (Fig. 4A; Stoodley et al., 2018). In each case, DREADD expression was observed in the targeted nucleus and corresponding parasagittal regions of overlying Purkinje cells (Fig. 4B, Supp. Fig. 4A). Some labelling was also seen in a subset of DCN neurons that project to the contralateral inferior olive independent of which nucleus was targeted. We observed no expression or axonal labelling in pre-cerebellar pontine nuclei or any downstream premotor sites. The pattern of Purkinje cell labeling for each injection was generally well-restricted mediolaterally, with some partial overlap due to spillover around the injection site, while anteroposterior spread was broader (Fig. 4B; Supp. Fig. 4A).

**Figure 4.**
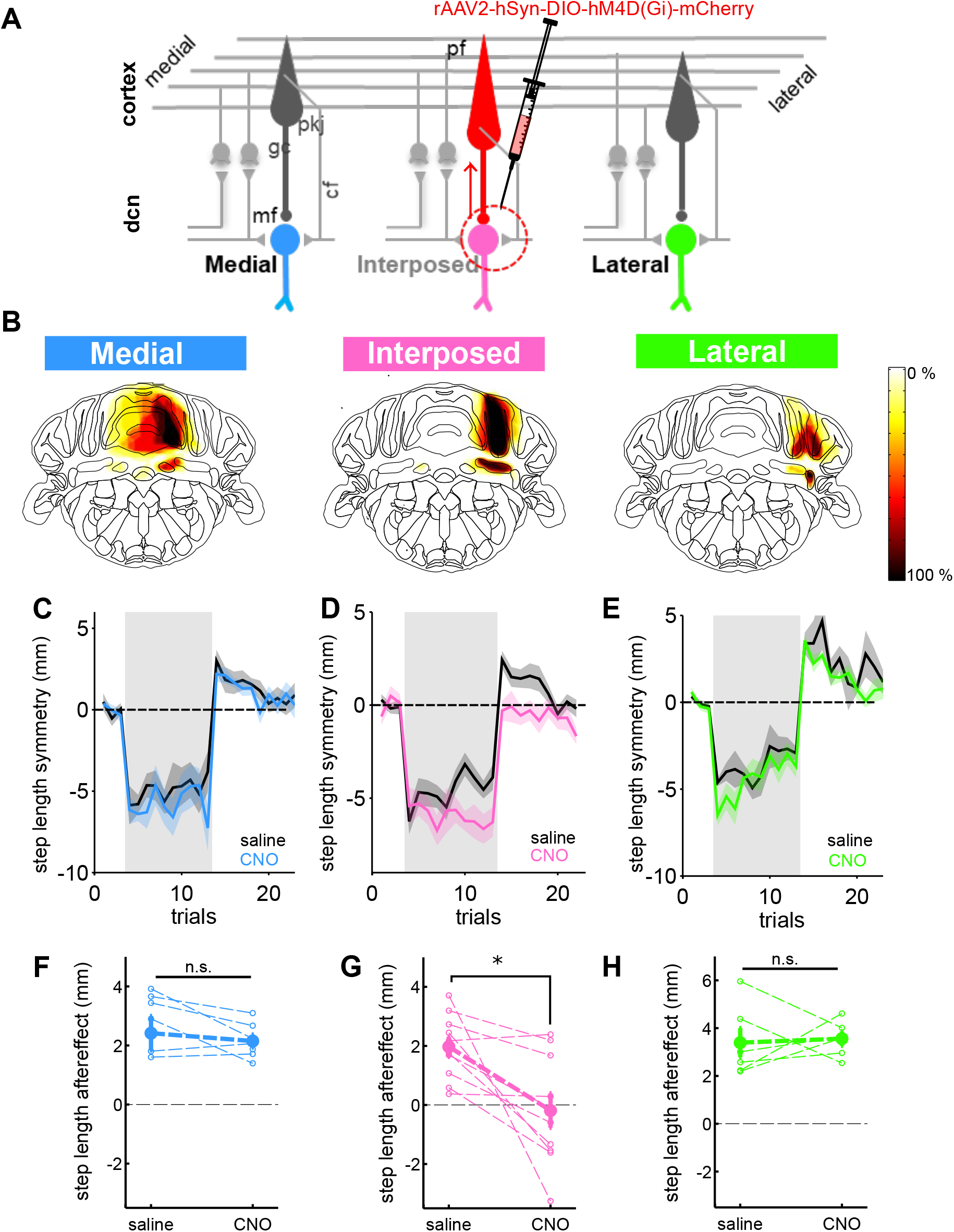
Locomotor adaptation requires projections to the interposed cerebellar nucleus. A) Cerebellar circuit diagram depicting retrograde approach for targeting inhibitory DREADDs to Purkinje cells terminating in each of the three cerebellar nuclei. Cerebellar structures and cell types are indicated with abbreviations: dcn: Deep cerebellar nuclei; mf: mossy fibers; gc: granule cells; pkj: Purkinje cells; cf: climbing fibers. B) Average virus expression area for medial, interposed and lateral nucleus injections, overlaid on mouse brain atlas. Colorbar represents percent of mice with expression in a particular zone. (C-E) Average step length adaptation (± s.e.m.) curves for medial (C, blue; N = 7), interposed (D, pink; N = 10), and lateral (E, green; N = 6) nucleus injections, following administration of saline (black) or CNO (colors). (F-H) Average step length after-effect sizes (first 2 washout trials) for saline and CNO conditions for medial (F), interposed (G) and lateral (H) nuclei. Individual animals are represented with thin, dashed lines, and open circles, averages with closed symbols. Significance was assessed with paired *t*-tests.

Injections into the right medial nucleus labelled Purkinje cells in the vermis of lobules 3-6, with partial spillover to most medial parts of interposed and paravermal cortex (Fig. 4B; Supp. Fig. 4A). Interposed nucleus injections labeled Purkinje cells in paravermal zones along the primary fissure, throughout lobules 2-6 (Fig. 4B, Supp. Fig. 4A). We observed expression throughout the entirety of the interposed nucleus, including anterior, posterior and dorsolateral subzones. Injections targeting the lateral nucleus labelled Purkinje cells in lateral simplex, and crus 1-2 (Fig. 4B, Supp. Fig. 4A).

Importantly, neither systemic CNO nor repeat adaptations interfered with adaptation; non-DREADD expressing animals adapted similarly with saline and upon subsequent administration of CNO (Supp. Fig. 5A,B; see also Supp. Fig. 2D; saline v CNO after-effect for step length: *t*_(7)_ = −.87, *p* = 0.41).

Two weeks after virus injection, mice were tested in split-belt adaptation experiments. We compared the adaptation exhibited by individual DREADD-expressing mice after administering saline vs. clozapine-N-oxide (CNO; 10 mg/kg) intraperitoneally (Fig 4C-H). Mice underwent adaptation protocols in which the fast belt was ipsilateral to the virus injection (ipsi-fast).

In mice with medial nucleus injections we observed no difference in step length adaptation between saline and CNO conditions (aftereffect, CNO v saline: *t*_(6)_ = .53, p = .55; Fig. 4C,F). Similar to the medial nucleus injections, we also found no differences between saline and CNO conditions when targeting the lateral nucleus (Fig. 4E,H, step length after-effect ipsi-fast: *t*_(5)_ = −.24, *p* = .82). In contrast, for interposed injected animals, administration of CNO impaired step length adaptation (Fig. 4D,G; aftereffect, CNO v saline: *t*_(9)_ = 3.24, *p* = .01). Importantly, DREADD activation in these animals did not cause overt ataxia and we found no differences in baseline (tied belt) step length symmetry between saline and CNO conditions (*t*_(9)_ = −.86, *p* = .41) or differences in stride length scaling in early split (*t*_(9)_ = −.93, *p* = .38), indicating little perturbation of baseline locomotion (Supp. Movie 2).

These cell-type specific chemogenetic experiments suggested a specific role for paravermal Purkinje cells in split-belt adaptation. As an independent test of this hypothesis, and since the retrograde virus also showed some expression in nucleo-olivary projection neurons, we next injected pan-neuronal DREADDs directly in the paravermal cerebellar cortex. Direct manipulation of this region of cerebellar cortex was sufficient to impair step length adaptation (Supp. Fig. 5, C-E, step length aftereffect, CNO v saline: *t*_(3)_ = 3.64, *p* = .036).

Finally, in separate groups of animals, we tested the effects of unilateral DREADD activations with the fast belt contralateral to the injection site (contra-fast). As with the ipsi- fast condition, step length adaptation was only impaired following administration of CNO in interposed-injected mice (Supp. Fig. 4B-G; contra-fast step length after-effect, CNO v saline: *medial:* t (4) = −.65, p = .62; *interposed* : *t*_(6)_ = 3.05 *p* = .02; *lateral: t*_(2)_ = .68, *p* = .56).

### What is being learned?

Because step length is a global measure that compares the movements of two limbs relative to each other, it obscures information about how each limb contributes to adaptation. In order to elucidate potential differential contributions of adaptation of individual limbs to overall locomotor adaptation, we analyzed changes in spatial and temporal parameters for each of the four limbs.

Although step length adaptation in hind limbs was modest (Supp. Fig. 1B,C; Supp. Fig. 6A), breaking it down into spatial and temporal components in the long experiment revealed that hind limbs exhibited significant spatial adaptation (Supp. Fig. 6B; *F_(3, 16)_* = 299.23, *p* < .001; change over split: *t*_(16)_ = −3.22, *p* = .005; after-effect: *t*_(16)_ = −4.45, *p* = .001) that proceeded more slowly than front limb spatial adaptation (coo: λ = 1804 strides). In contrast, the hind limbs did not exhibit adaptation of temporal symmetry (Supp. Fig. 6C; over the split-belt period: *t*_(16)_ = .81, *p* = .42; after-effects: *t*_(16)_ = 1.71, *p* = .1). Thus, the small degree of step length adaptation observed in hind paws comes purely from spatial, and not temporal, adaptation.

We next examined the contributions of each of the four individual limbs to spatial adaptation. In early split, both limbs on the fast belt oscillated on an axis further back on the body than the limbs on the slow belt (Fig. 5A). Over split-belt trials, all four limbs adapted; fast limbs shifted forward and slow limbs backward on the body with a magnitude that was similar across all paws, with after-effects in the opposite directions (Fig. 5 A-C; after-effect magnitude comparison across four paws: *F_(3, 16)_* = 1.87, *p* = .15).

**Figure 5.**
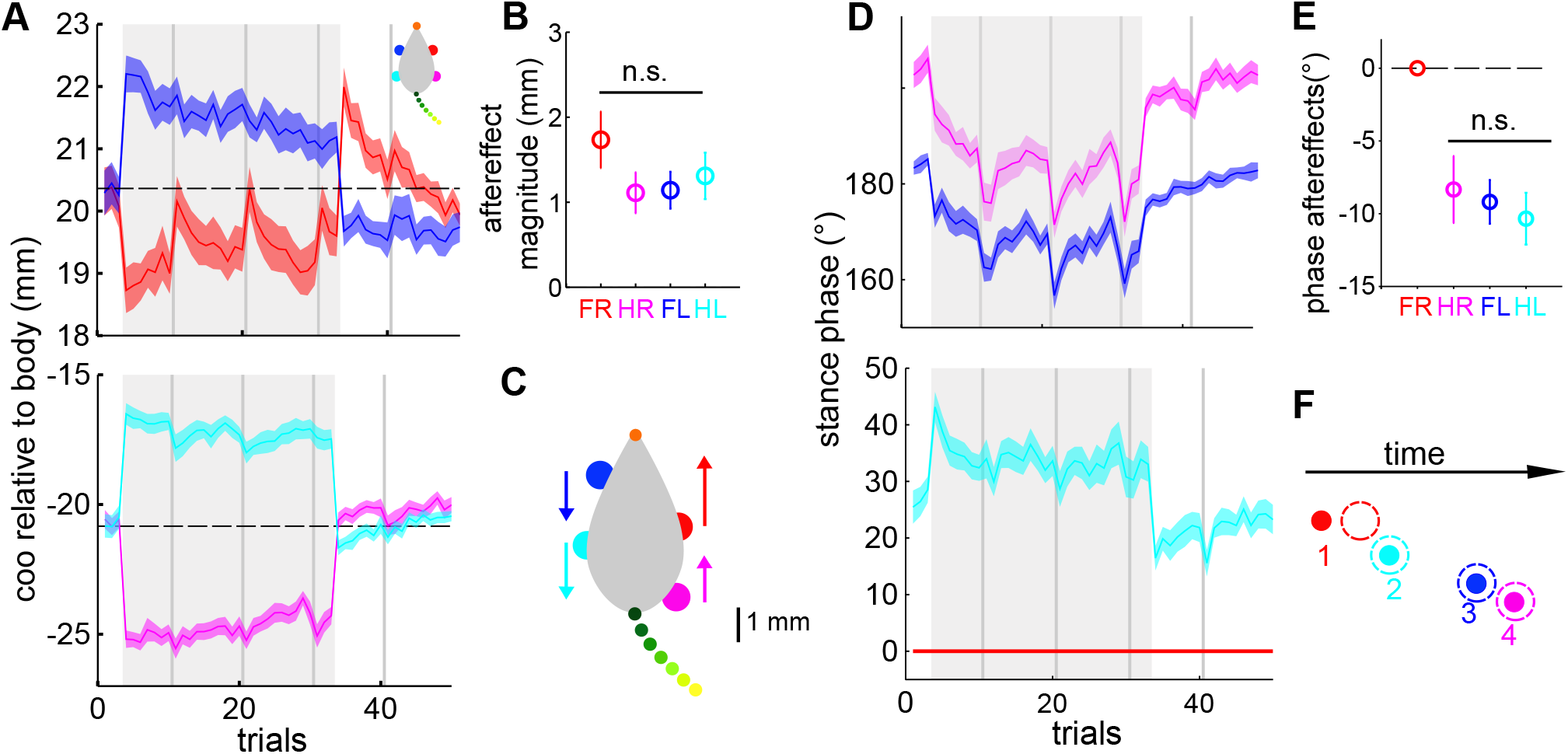
Differential contributions of the four individual limbs to spatiotemporal adaptation. A-C) Spatial adaptation. A) Average (n = 17) center of oscillation for each limb over an experiment with 30 minutes of split-belt walking. The color code for the four limbs is represented in the inset at top right. B) Average spatial aftereffect magnitude for each limb. There was no significant difference in aftereffects compared across the four limbs. C) Schematic of early split and learned changes for the spatial parameter. In early split, limbs on the fast belt (red, magenta) oscillate around a more caudal axis relative to the body than limbs on the slow belt (blue, cyan). Over adaptation, fast limbs shift their center of oscillation rostrally while slow limbs shift caudally, as indicated by the arrows. Scale bar represents the magnitude of spatial shift for each limb (mouse body not to scale). D-F) Temporal adaptation. D) Average stance phase of each limb relative to front right (fast) limb over the course of the same experiment. E) Average after-effect for each limb, relative to the front right (fast) reference limb. F) Illustration of one possible configuration of learned changes for the temporal parameter. The relative timing of stance onset for each limb during early split is represented with solid circles, the open, dashed circles indicate the shifts over the adaptation period. The shift of the front, fast paw relative to the other three restores the normal diagonal limb alternation pattern.

The need for a stride-cycle temporal reference (as opposed to a body-centric spatial reference) complicates resolving the individual limb contributions to temporal adaptation. We examined temporal adaptation for individual limbs by measuring stance phasing for all paws relative to the front fast limb (Fig. 5 D-F). Mice normally trot across speeds, with diagonal limbs entering stance together and out of phase with the opposite diagonal pair (Machado, Darmohray et al. 2015; Fig. 5D, F, baseline). In early split, the normal pattern of inter-limb coordination is disrupted: relative to the front fast reference paw, its diagonal partner touches down later, while both paws in the opposite diagonal pair touch down earlier (Fig. 5D, illustrated in Fig. 5F, baseline). Over the split-belt period, all three paws advanced their stance onset relative to the front, fast reference paw, resulting in a recovery of the diagonal pairing (Fig. 5E, after-effect comparison across non-ref paws: *F_(2, 16)_* = .36, *p* = .7). Since the hind limbs did not show temporal adaptation relative to each other, this implies that either the front fast paw adapted relative to the other three, or vice versa. Similar results were obtained when the belt on the left side was run fast, indicating an effect limb speed, rather than handedness (after-effect comparison across non-ref paws: *F*_(4_, _2)_ = 1.31, *p* = .32).

### Distinct cerebellar contributions to spatial and temporal adaptation

The results of Fig. 5 suggest that while spatial adaptation involves changes across all four limbs, the front, fast limb might make a unique contribution to temporal adaptation. We therefore wondered how spatial and temporal learning would be lateralized in the cerebellum. In particular, given that the cerebellar control of movement is predominantly ipsilateral, we hypothesized that chemogenetic manipulations in interpositus might interfere with temporal adaptation only when injections were made ipsilateral to the fast belt. Similarly, since spatial adaptation appears to involve limbs on both sides of the body, we hypothesized that it might be affected by unilateral chemogenetic manipulation in both the ipsi- and contra-fast conditions.

We addressed this question by separately analyzing the effects of CNO on spatial and temporal adaptation in the interposed injected animals under conditions in which the belts were run fast either ipsi- or contra-fast relative to the injection (Fig. 6A). As predicted, spatial adaptation was impaired in both the ipsi-fast (coo, CNO v saline aftereffect: *t*_(9)_ = 3.32, *p* = .009) and contra-fast conditions (Fig. 6 C,D, CNO v saline aftereffect: *t*_(8)_ = 2.81, *p* = .03). In contrast, temporal adaptation was impaired with CNO administration only in the ipsi-fast condition (Fig. 6E, left; double support symmetry, CNO v saline aftereffect: *t*_(9)_ = 2.63, *p* = .02). Temporal adaptation was not different from control in the contra-fast condition (Fig 6E, right; *t*_(6)_ = −1.49 *p* = .2). Interestingly, when spatial adaptation was selectively impaired (in the contra-fast condition), we did not observe a compensatory over-adaptation of temporal symmetry, that could have helped equalize step length (Fig. 2G). Thus, the separate time courses of spatial and temporal adaptation, together with their differential lateralization both behaviorally and in the cerebellum, suggest that locomotor learning is achieved through independent spatial and temporal learning processes, with different circuit implementations.

**Figure 6.**
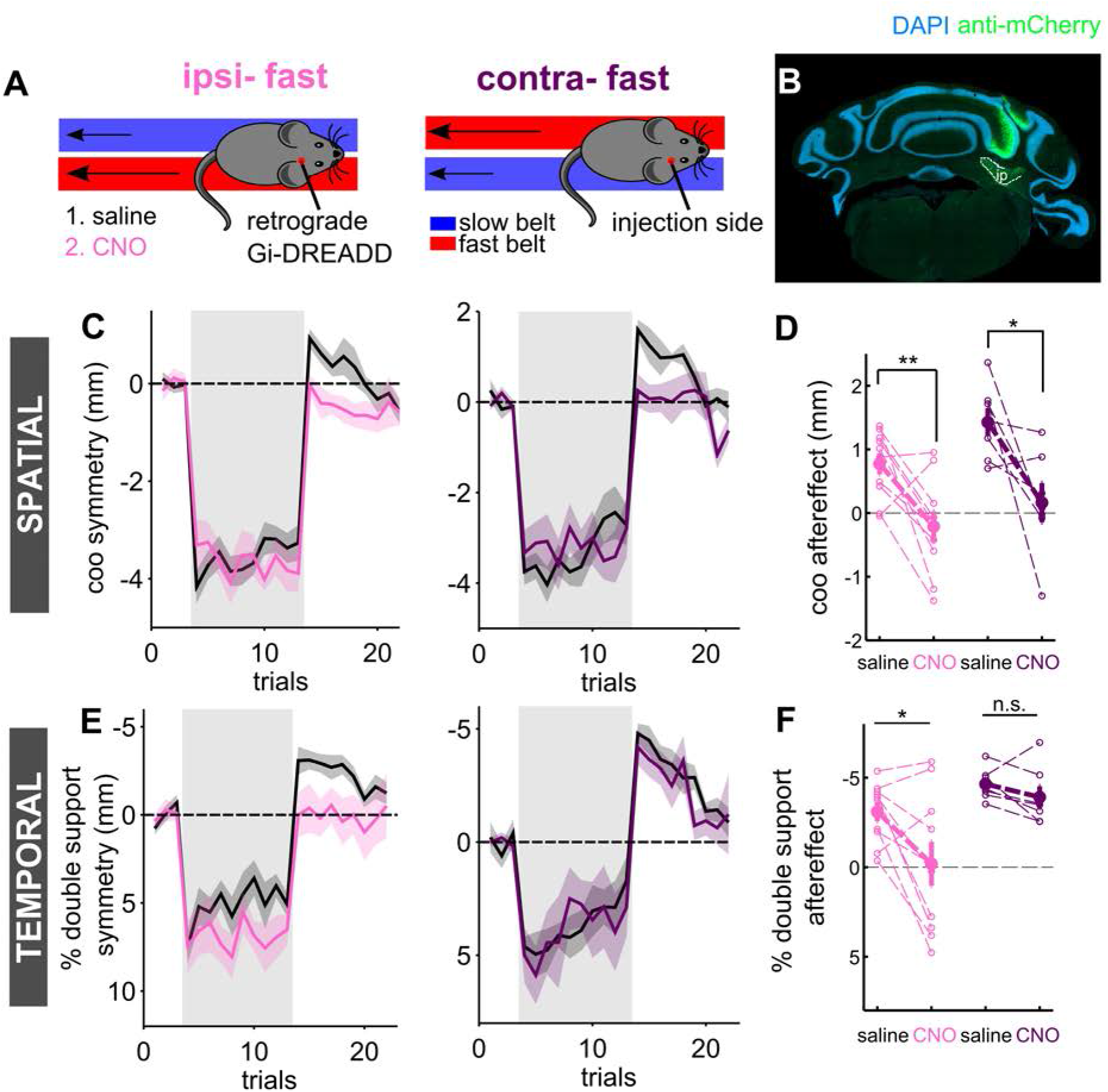
Differential lateralization of spatial and temporal learning in the cerebellum. A) Experimental protocol for assessing lateralization of split-belt learning. Animals with unilateral cerebellar injections underwent adaptation protocols where the fast belt was either ipsilateral (ipsi-fast, right) or contralateral (contra-fast, left) to the cerebellar injection site. Mice in each condition were run first with intra-peritoneal of saline, and subsequently with intra-peritoneal injections of CNO. B) Example cerebellar section from a mouse with unilateral injection of retrograde DREADDs in the interposed nucleus. The borders of the interposed nucleus (ip) are indicated by a white, dashed line. Sample was stained with DAPI (blue) and anti-mCherry (for DREADD expression). C) Average front limb center of oscillation (spatial) adaptation curves (± SEM) for ipsi-fast (left, pink) and contra-fast (right, purple) conditions. In each plot, black curves represent mice under saline conditions; colors represent CNO. D) Average center of oscillation after-effect magnitudes (first 2 washout trials) for ipsi- and contra-fast conditions, with saline and CNO. Individual animals are shown with thin, dashed lines. E) Average double support (temporal) adaptation curves (± SEM) for ipsi- and contra-fast conditions, plotted as in C. F) Average center of oscillation aftereffects (first 2 washout trials) for ipsi- and contra-fast conditions, with saline and CNO. Significance was assessed with paired *t*-tests.

## Discussion

We developed a transparent split-belt treadmill for mice that provides high resolution readouts of locomotor behavior using a noninvasive, markerless tracking system (LocoMouse; Machado, Darmohray et al., 2015). We found that mice learn to adapt their locomotor patterns to regain symmetry in response to external perturbations in which treadmill belts under each side of the body move at different speeds. Mouse split-belt locomotor adaptation is cerebellum-dependent, specific to measures of interlimb coordination, and has spatial and temporal components with distinct time courses. We identified a subregion of the cerebellar cortex whose Purkinje cells terminate in the interposed nucleus that is required for this form of locomotor learning. Consistent with predictions from the analysis of learned changes in spatial and temporal gait components, unilateral chemogenetic manipulations in this region differentially impaired spatial and temporal adaptation. These findings provide insights into how the movements of four independent limbs are calibrated and coordinated by cerebellar circuits during locomotion.

### Mouse split-belt locomotor adaptation

Previous studies have demonstrated locomotor adaptation on a split-belt treadmill in cats (Yanagihara and Kondo, 1996; Yanagihara et al., 1993) and humans, where it has been extensively studied and is used as a rehabilitation therapy (Reisman et al., 2005, 2007, 2010). Here, we implemented split-belt adaptation for the first time in mice, in order to probe neural circuit mechanisms underlying this form of motor adaptation. Our results show that mice learn in a way that is remarkably similar to humans. Like human learning, mouse locomotor adaptation is specific to interlimb coordination and has spatial and temporal components that adapt at different rates (Malone et al., 2011, 2012; Reisman et al., 2005). Further, split-belt adaptation in mice is dependent on an intact cerebellum, but is insensitive to large lesions of cerebral cortex (Fig. 3). Finally, it is also exhibited high levels of retention and did not generalize to other sensorimotor contexts (ie, overground walking), features that it shares with not only human split-belt treadmill adaptation (Reisman et al., 2009; Torres-Oviedo and Bastian, 2010), but also various forms of cerebellum-dependent learning (Bodznick et al., 1999; Kahlon and Lisberger, 1996; Miles and Eighmy, 1980; Straube et al., 1997).

Overall, the striking similarities between mouse and human learning suggest that the mechanisms underlying locomotor adaptation are highly conserved, opening up the possibility of using genetic tools in mice to elucidate the neural mechanisms underlying locomotor adaptation across species. The similarities between mouse and human locomotor adaptation are perhaps surprising given the many differences in quadrupedal vs bipedal locomotion. In fact, the only difference we observed between mouse and human locomotor adaptation was a lack of savings in mice (Malone et al., 2011). Notably, a lack of savings has been previously observed for other forms of cerebellum-dependent motor adaptation (Huang et al., 2011; Kahlon and Lisberger, 1996; Miles and Eighmy, 1980; Miles and Lisberger, 1981; Straube et al., 1997). One intriguing possibility is that the presence of savings in human split-belt adaptation may be accounted for by the enhanced contribution of cerebral cortex to locomotion in humans and higher vertebrates (Massion, 1967).

### Locomotor adaptation is specific to learned changes in interlimb coordination

While basic gait patterns can be generated within the spinal cord, they are influenced by sensory input and supraspinal centers that work together to coordinate and modulate locomotion. In the split-belt experiments conducted here, animals exhibit multiple responses to the externally-imposed perturbations of belt symmetry. There are immediate changes in stride length, for example, that persist throughout the split-belt period; stride symmetry is regained only when the two belt speeds return to symmetrical baseline levels. These adjustments, like the ability of spinal cats to respond to changes in treadmill speed by transitioning from one gait to another, represent reactive changes to belt speed and do not require learning (Forssberg et al., 1980; Reisman et al., 2005; Whelan, 1996).

Changes to locomotor patterns that are learned and stored in the brain, however, are typically characterized by gradual changes throughout the split-belt period and are accompanied by after-effects once treadmill belt symmetry is restored. Similar to human cerebellar patients (Morton and Bastian, 2006), while *pcd* and *reeler* mutant mice were able to rapidly adjust their stride lengths (a reactive, individual limb parameter) in response to changes in belt speeds, they showed absolutely no evidence of learning, even after prolonged exposure to split-belt walking. The complete abolition of learning in these mutants, both of which present with cerebellar malformation from an early age (Lalonde and Strazielle, 2007), is perhaps surprising. First, partial learning is generally observed in human focal cerebellar lesion patients (Vasudevan et al., 2011). Second, these mutations are chronic, and there is, in theory, plenty of time for compensatory mechanisms to potentially overcome the cerebellar degeneration. Thus, this finding demonstrates a remarkably specific role for the cerebellum in learned changes in interlimb coordination during split-belt walking, which nicely parallels our previous finding that the visible gait ataxia of *pcd* mice is specific to impairments in interlimb coordination, while individual limb gait parameters (like stride length) remained intact (Machado, Darmohray et al., 2015). These results also contrast with the intact adaptation we observed following widespread lesions of somatomotor cortex. While we cannot rule out that a possible role of somatomotor cortex may have been masked by compensatory mechanisms that arose during the days following the lesion, we note the comparative inability of any such putative mechanism to compensate for the chronic cerebellar damage in the mutants.

With prolonged split-belt exposure, front limbs nearly completely recovered baseline symmetry for both temporal and spatial adaptation. In contrast, hindlimb adaptation was incomplete, purely spatial, and occurred at a slower rate than front limbs. One possibility is that the partial spatial adaptation observed in hindlimbs partially reflects adjustments to match the new, learned patterns of front limb motion, rather than representing learning itself. Consistent with this idea, the cerebellum exerts its influence on locomotion mainly via projections to midbrain and brainstem premotor nuclei (Liang et al., 2011), many of which preferentially target forelimb motor neurons (Esposito et al., 2014). Front-hind limb matching could then be achieved through long-range cross-segmental spinal communication or other reflexive spinal mechanisms that represent only secondary consequences of cerebellar learning (Forssberg et al., 1980; Machado, Darmohray et al., 2015; Ruder et al., 2016).

### Cerebellar circuits for locomotor adaptation

We used a chemogenetic approach to identify a population of Purkinje cells in paravermal cortex projecting to the interposed nucleus that are required for split-belt treadmill adaptation. Because of the lack of savings for this form of learning in mice, we were able to use within-animal controls to demonstrate that CNO affected locomotor adaptation only when inhibitory DREADDS were expressed in this cerebellar subregion (Roth, 2016, 2017). These results were remarkably specific; CNO had no effect on locomotor learning when the same viral strategy was used to target DREADDs to nearby regions. Moreover, the same results were observed regardless of the viral strategy used to deliver the DREADDS to these Purkinje cells (Supp. Fig. 5).

The ability to localize split-belt adaptation to such a relatively small cerebellar subregion is perhaps surprising in the context of the widespread representation of sensorimotor signals associated with locomotion across the cerebellum (Armstrong and Edgley, 1984; Armstrong et al., 1988; Edgley and Lidierth, 1988; Marple-Horvat and Criado, 1999; Ozden et al., 2012; Powell et al., 2011; Sarnaik and Raman, 2018; Udo et al., 1981). Indeed, many cerebellar regions contribute to locomotion in various ways. Both medial and interposed cerebellar nuclei receive sensory and state information from spinal locomotor circuits, and activity in Purkinje cells projecting to them is modulated by the step cycle (Armstrong and Edgley, 1984; Edgley and Lidierth, 1988). Meanwhile, the lateral cerebellum has been implicated in the overriding of locomotor patterns, such as obstacle avoidance, although damage in this zone does not markedly affect normal gait (Aoki et al., 2013; Marple-Horvat and Criado, 1999).

However, anatomical and lesion studies, together with patient literature, have demonstrated a clear medial to lateral functional organization within the cerebellum (Chambers and Sprague, 1955; Morton and Bastian, 2007). For instance, the medial zone is highly interconnected with the vestibular system, and accordingly, midline damage disrupts balance and posture(Chambers and Sprague, 1955; Morton and Bastian, 2004; Takakusaki, 2017). Meanwhile, the lateral cerebellum largely targets thalamocortical pathways, and the intact adaptation we observed with DREADD manipulations here is consistent with the results of the widespread aspirations of sensorimotor regions of the cerebral cortex (Fig. 3) and with the idea that split-belt adaptation is mediated by more subcortical anatomical pathways (Reisman et al., 2007) . In contrast, damage to intermediate regions cause more subtle deficits that suggest that it is particularly important for precise limb placement and interlimb coordination (Chambers and Sprague, 1955; Udo et al., 1979, 1980) – exactly the parameters that undergo learned changes during split-belt adaptation.

Several aspects of our data point to distinct circuit mechanisms underlying the spatial and temporal components of the locomotor learning described here (Choi et al., 2009; Malone and Bastian, 2010; Malone et al., 2011; Torres-Oviedo and Bastian, 2010; Vasudevan et al., 2011). First, as in humans, we found that temporal adaptation proceeds faster than spatial learning. Second, spatial and temporal learning involve differential contributions from the four limbs. While all four limbs appear to undergo spatial adaptation, our analyses suggest that in normal temporal adaptation, the front fast limb adjusts its timing relative to the other three limbs. Third, consistent with the interlimb coordination analyses, chemogenetic manipulations affected spatial adaptation bilaterally, but only interfered with temporal adaptation when applied ipsilateral to the fast belt. Finally, manipulations that specifically blocked spatial adaptation did not result in a compensatory over-adaptation of temporal parameters, that might be expected to occur if step-length itself were being optimized (Fig. 2G). While the precise error signal or signals that are being corrected for in locomotor adaptation are not known (Finley et al., 2013), the fact that spatial and temporal learning appear to occur independently may suggest that they represent adaptive responses to separate error signals.

The distinct circuitry for spatial and temporal components of locomotor learning suggested here are particularly interesting in light of previous experiments demonstrating a role for the cerebellum in precise timing (Ivry, 1996; Ivry and Spencer, 2004; Medina and Mauk, 2000),as well as studies that have suggested distinct circuit mechanisms for timing vs. amplitude in other cerebellar learning tasks (Medina et al., 2000; Raymond et al., 1996). It will be interesting to further map the downstream connectivity of cerebellar interposed projections to better understand split-belt adaptation and coordination across the limbs and in space and time. It is possible that space and time will be further segregated in specific neural populations within the cerebellar cortex, interposed nucleus and/or downstream circuitry.

### Conclusion

Locomotor adaptation on a split-belt treadmill represents a new form of motor learning in mice that provides good experimental control in the context of a natural behavior that is highly conserved across vertebrates. It is a rapid form of cerebellum-dependent learning that can be measured non-invasively in freely-moving mice with minimal previous training. The ability to distinguish adjustments in individual limb movements (ie stride length) from learned changes in interlimb coordination (like step length) can be exploited in order to assess specific deficits in locomotor coordination and learning in various mouse models. Moreover, it offers a powerful opportunity to combine quantitative behavioral analysis with genetic tools for circuit dissection to gain insight into neural mechanisms for motor learning.

## Acknowledgements

We thank Tracy Pritchett for maintenance of mouse lines and help with histology and imaging, and Joao Fayad for optimization and maintenance of the LocoMouse tracker for the split-belt treadmill. Lorenza Calcaterra, Olivia Carmo, and Marta Maciel assisted with data acquisition and Inês Vaz provided surgical support. We are grateful to the members of Carey lab and the Champalimaud Neuroscience Program for helpful discussion.

This work was supported by a Howard Hughes Medical Institute International Early Career Scientist Grant #55007413 (to MRC), European Research Council Starting Grant #640093 (to MRC), and fellowships from the Portuguese Fundação para a Ciência e a Tecnologia SFRH/BD/86265/2012 (to DD), SFRH/BD/52450/2013(to JJ), and #BPD109659/2015 (to HGM).

## Author contributions

Conceptualization, DD and MRC; Methodology, DD and HGM; Conducting experiments, DD and JJ; Data analysis, DD, JJ, MRC; Writing – Original Draft, DD and MRC; Writing – Review and Editing, DD, HGM, JJ, MRC; Supervision, MRC; Project Administration, MRC; Funding Acquisition, MRC

## Methods

### Animals

All procedures were reviewed and performed in accordance with the Champalimaud Centre for the Unknown Ethics Committee guidelines and approved by the Portuguese Direcção Geral de Veterinária. Mice were housed in institutional standard cages (5 animals per cage) on a reversed 12-h light/12-h dark cycle with *ad libitum* access to water and food. Heterozygous Purkinje cell degeneration and reeler mice on a C57BL/6 background were obtained from Jackson labs (pcd: #0537, B6.BRAgtpbp1pcd/J; reeler: #000235,B6C3Fe *a/a-Relnrl/J*). L7-cre mice used in DCN DREADDs manipulations were also obtained from Jackson labs (# 004146, Tg(Pcp2-cre)1Amc/J).

### Experimental setup

We developed a transparent split-belt treadmill for mice that provides high resolution readouts of locomotor behavior using the previously described noninvasive, markerless LocoMouse tracking system (Machado, Darmohray et al., 2015; https://github.com/careylab/LocoMouse). Briefly, mice were filmed while walking inside a narrow corridor (30 x 4 cm). A 45° angled mirror below the setup allowed for simultaneous collection of side and bottom views (Machado, Darmohray et al., 2015). Lighting consisted of a matrix of LEDs that emitted cool white light positioned to maximize contrast and reduce reflection. Transparent mylar belts were driven using two DC motors with high-resolution encoders and an Escon 50/5 motor controller (Maxon). Images were acquired at 330 fps using a monochrome FireWire camera (Pike F-032 B/C, Allied Vision Technologies, https://www.alliedvision.com) and custom-software written in Labview.

### Split-belt adaptation protocols

Animals were handled and acclimated to the treadmill prior to starting experiments. Treadmill acclimation consisted of 3-4 daily sessions of tied belt walking until mice walked without turns, maintained a regular position, and were able to keep up with the range of belt speeds used in the adaptation protocol. Experimental mice were freely moving and ran without reinforcement. Although there was no barrier between belts, the narrow belt widths (2 cm) prevent prolonged paw crosses to belts on the opposite side. We ensured adherence to imposed treadmill speeds by monitoring the average paw speeds during the stance period.

All split-belt locomotor adaptation experiments consisted of ‘baseline’ tied, split-belt, and ‘washout’ tied belt trials. All trials were one minute in duration, with brief periods in which the motors were off, in between trials. Tied-belt baseline trials were at an intermediate speed between slow and fast sides (generally 0.275 m/s). Split-belt trial speeds were generally at a 2.14:1 ratio: 0.175 m/s (slow) and 0.375 m/s (fast). For PCD and reeler mutant mice, the overall belt speeds were reduced but a similar ratio between fast and slow was maintained (tied: 0.225 m/s; slow: 0.125 m/s fast: 0.325 m/s).

Our extensive experience, including but not limited to the data presented in Supp. Fig. 2 and Figs. 4 and 6, clearly indicates that there is no difference in learning following repeated adaptation protocols in individual animals, as long as learning is washed out completely in between experiments. Therefore, in all experiments involving surgical intervention, we used within-animal controls, to take advantage of the ability to directly compare learning under different experimental conditions in the same mice.

### LocoMouse tracking and gait analysis

Paw, nose and tail tracks were obtained using LocoMouse tracker: an open-source, 3D markerless tracking algorithm described previously (https://github.com/careylab/LocoMouse; Machado, Darmohray et al, 2015). Periods of inactivity and/or egregious tracking failures were removed via an algorithm that excluded data with physically impossible track configurations (e.g, left-right or front-hind paw inversions) and those where no strides were detected. On average, 70.5% of tracks were retained, resulting in an average of 179.4 +/−36.96 (*SD*) strides each minute. Tracks were then subdivided into stride cycles by automatic detection of swing and stance points using a simple peak detection algorithm for the stride-wise computation of all gait parameters (Machado, Darmohray et al. 2015).

Individual strides were defined from stance onset to subsequent stance onset. Measures of symmetry are always reported as the fast minus the slow side for homologous limbs (contralateral limbs sharing a girdle; e.g, front left - front right; hind left - hind right) and were computed on the average trial values for each individual limb. In all reports of symmetry, measures were baseline subtracted using the average values for baseline tied-belt trials. Individual limb and interlimb coordination parameters were calculated as follows:

*Step length:* displacement of one limb relative to its (contralateral) homolog at stance onset
*Stride length:* total displacement of an individual limb from swing onset to stance onset
*Center of oscillation:* midpoint between swing and stance position (along rostrocaudal axis) of an individual limb relative to the body center
*% double support* - percentage of the stride cycle duration in which two (contralateral) homologous limbs are simultaneously in stance
*stance phasing:* relative timing of limb stance onsets within the stride cycle of a reference paw. Calculated as: (stance time - stance time_ref._)/stride duration_ref._

### Statistics

All statistical analyses were done using Matlab. For basic split-belt adaptation experiments (Figs 1-2), we used repeated measures analysis of variance (ANOVA) to test for significant effects of experimental phase (baseline, early split, late split, early washout) on the symmetry of each gait parameter (either stride length, step length, coo, or % ds). We defined early and late split as the first and last trial of the split-belt period, respectively, and early after-effect as the first 1-2 washout tied-belt trials. Follow-up for significant *F* tests were conducted using repeated measures *t*-tests in which we compared one or more of the following: baseline to early split, early to late split, and aftereffect to baseline. For mutant data and DREADD manipulations, we used the early washout period to quantify learning and compared experimental vs. control conditions with *t*-tests (paired, when applicable). We used a significance cutoff of p < 0.05 for all statistics. Statistical significance in plots is indicated by * for p < 0.05, ** for p < 0.01; and *** for p < .001.

### Simulated step lengths

Simulated stride cycles were generated by creating triangle waveforms with amplitudes, total stride durations, and durations for the stance/ swing phases of the stride cycle that were matched to average values for individual limb parameters in the early split-belt period (first split trial). Waveforms were then used to generate a predicted mapping of step length symmetry over many possible combinations of spatial and temporal shifts. To test the fit of the model to the experimental data, we matched experimental % double support and center of oscillation symmetry values with their closest simulation values and compared the actual step length values to the simulated step length symmetry for that spatiotemporal combination. Contributions of spatial and temporal parameters to step length symmetry were estimated by calculating slopes for unit shifts in each parameter vs step length, while holding the other constant.

### Primary motor and somatosensory lesions

Prior to aspirations, mice were habituated to the split-belt setup and underwent pre-lesion split-belt adaption to ascertain baseline learning, followed by complete washout. Somatomotor cortical lesions were performed by controlled vacuum aspirations via stereotaxic surgery with coordinates ⃞3.5 mm from midline; + 2.5 to −1.5 mm AP from Bregma; - 2.5 mm DV from brain surface. These coordinates were chosen to cover the full extent of forelimb primary motor and somatosensory cortices (Paxinos and Franklin, 2004). Buprenorphine (0.1 mg/kg) was administered intraperitoneally for postoperative analgesia. Mice were then housed in isolation and received injections of 0.01ml/g glucose and lactated Ringer’s solution for two consecutive days post-surgery. Animals were allowed to recover for before being re-habituated to the setup via a ten-trial tied-belt session. Split-belt locomotor adaptation took place five days following the lesions.

### DREADD experiments

Stereotactic surgery was performed under isoflurane anesthesia and buprenorphine (.1 mg/kg) was administered for postoperative analgesia. Intracranial injections were performed using a Nanoject II (Drummond) and 30-40 μm pulled and beveled glass pipettes. We injected retrograde, double floxed Gi-coupled hM4D DREADD (rAAV2 retro-hSyn-DIO-hM4D(Gi)-mCherry; addgene plasmid #44362, Tervo et al. 2016) into the deep cerebellar nuclei of L7-cre animals to target Purkinje cells terminating in each nucleus. For pan-neuronal cerebellar cortex injections (Supp. Fig 5), we injected AAV-hSyn-hM4D(Gi)-mcherry (addgene plasmid #50475, Armbruster et al., 2007). The following coordinates were used to target each of the deep cerebellar nuclei: medial: −6.24 AP, .5 ML, 2.2 DV; interposed: −6.24 AP, 1.6 ML, 2-2.3 DV; Lateral: −6 AP, 2.26 M, 2.2 DV. Anteroposterior (AP) and mediolateral (ML) coordinates are given relative to bregma. Dorsoventral (DV) are relative to the surface of the brain. The volume of all injections was between 100-200 nl and were on the right side. Mice used for these experiments were between 10-12 weeks old at time of injection. Split belt adaptation experiments were run two weeks later to allow time for virus expression. For CNO injections, we used dosages of 10 mg/kg and waited 30-40 minutes post intraperitoneal injection before starting the experiment. For all animals, learning was tested in the saline condition before the administration of CNO, and the ipsi and contra-fast conditions were tested in separate groups of animals. Mice were perfused with 4% paraformaldehyde 3-4 weeks post-injection.

## Supplemental information

**Supp. Figure 1.**
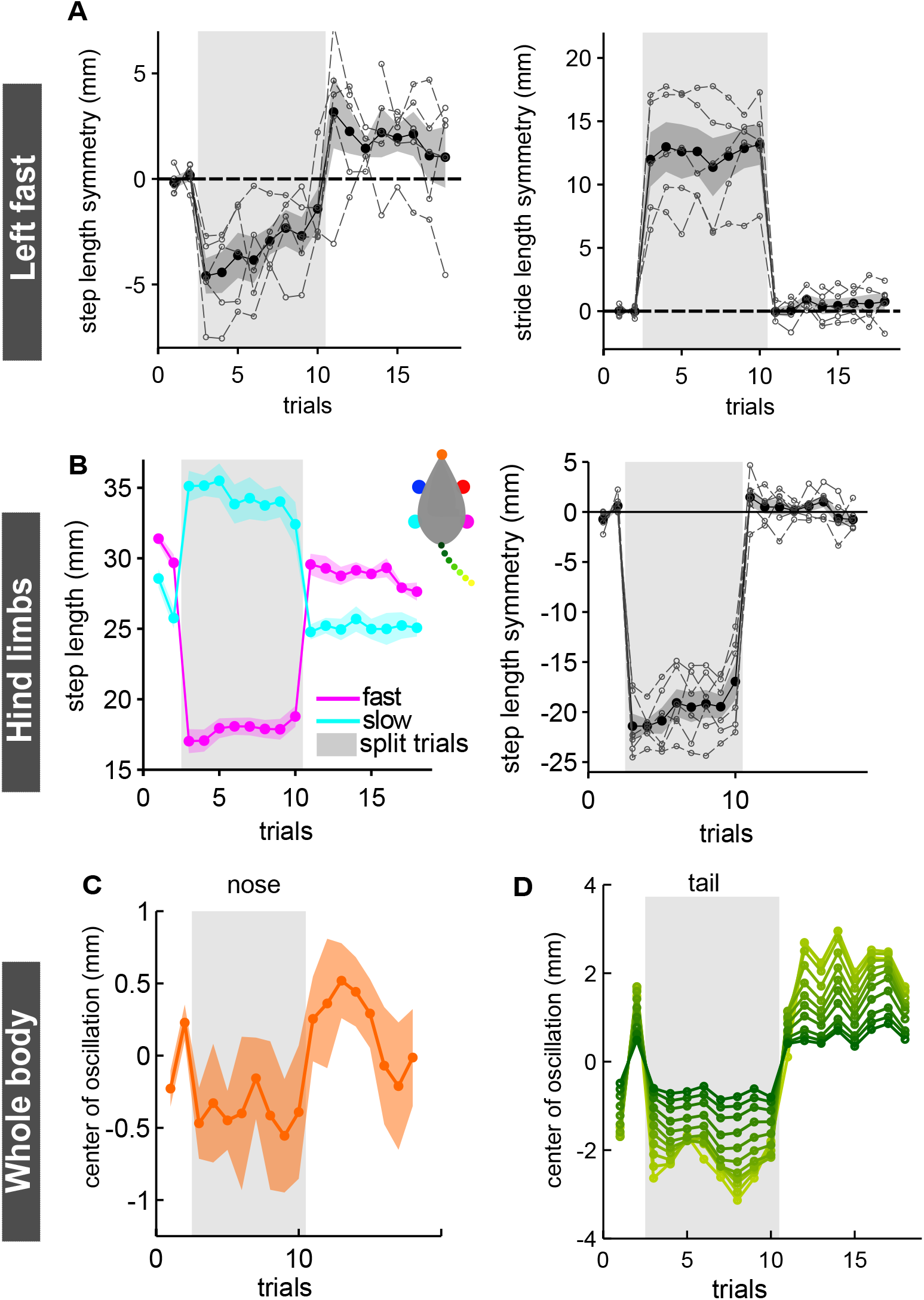
Left-fast, hindlimb, and whole-body movements during split-belt adaptation. A) left: Average (n=5) front limb step length symmetry (± SEM) for left fast experiment. right: Average front limb stride length symmetry (± SEM) for left fast experiment. Individual animal measures are represented with open symbols and dotted lines. Group average +/-SEM are indicated by closed symbols + shadow. B) left: Average (n=8) step length (± SEM) for fast (magenta) and slow (cyan) hind limbs over a single-session split-belt experiment. right: Average hind limb step length symmetry, plotted as in A. C) Average center of oscillation (± SEM) for nose in y (left-right) over split-belt trials of right fast experiment. Negative values indicate leftward movements D). Average center of oscillation of tail segments over split-belt trials of right fast experiment.

**Supp. Figure 2.**
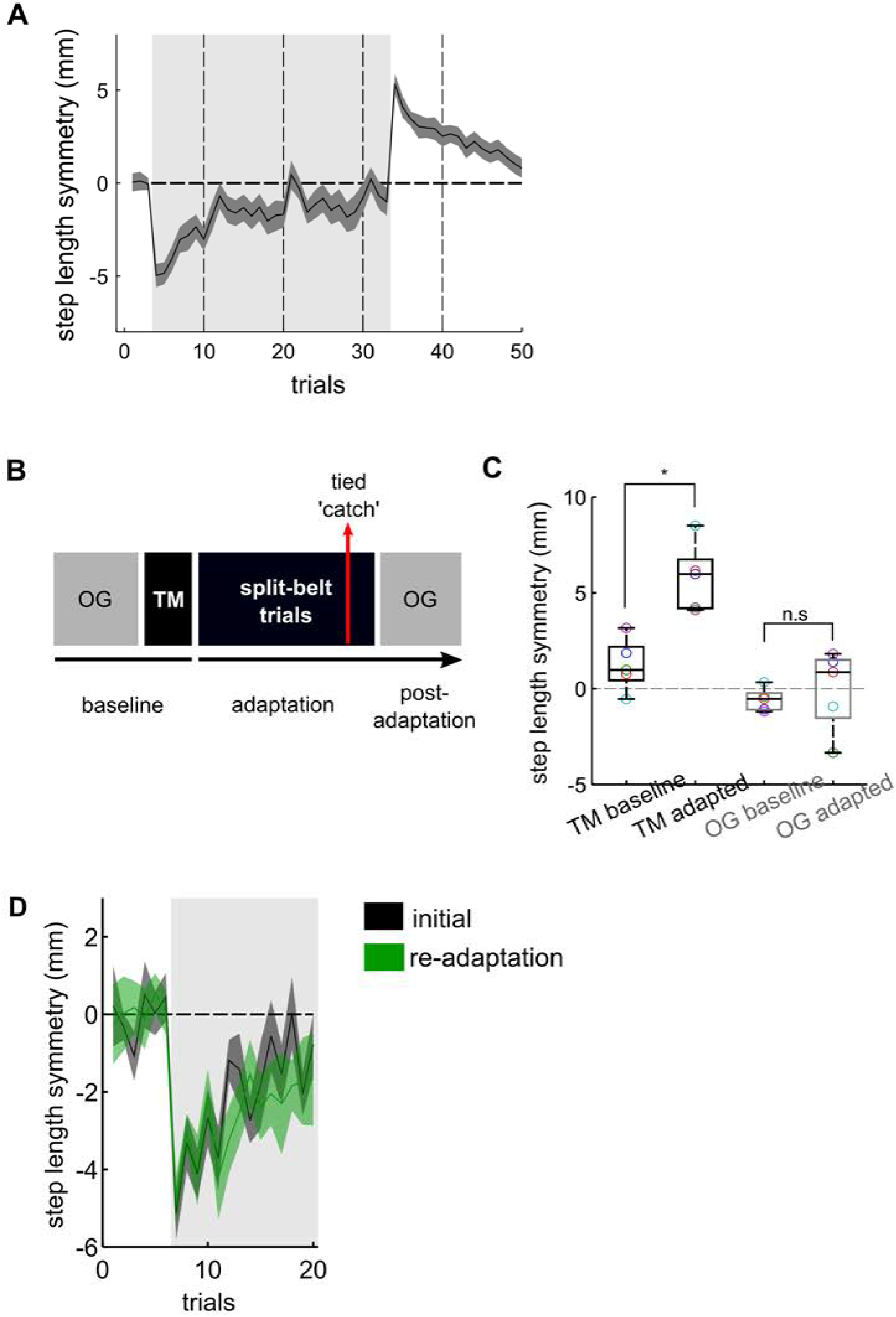
Mouse split-belt treadmill adaptation does not generalize across sensorimotor contexts and does not show savings. A) Average (n = 17) front limb step length symmetry over a 30-minute total split-belt period. Gray shaded patch indicates split-belt trials; darker vertical lines show session breaks. Shaded error represent SEM. B) Experimental protocol for assessing transfer of learning to a new walking context. Mice first walked overground to acquire baseline strides (OG baseline). Mice then underwent split-belt treadmill adaptation. A tied belt catch verified that mice showed significant aftereffects in symmetry relative to baseline on the treadmill (TM baseline v TM adapted,B). Finally, mice were moved to the overground setup in the TM-adapted state to test for the aftereffects in the new locomotor context (OG baseline v OG adapted,B). Overground strides were selected to match the tied belt speeds used in the adaptation protocol. C) Step length symmetry for two locomotion contexts: treadmill (TM; black) and overground (OG; gray); and adaptation states: baseline and adapted. Individual animals are shown as open circles for the four conditions. D) Average (n = 9) front limb step length symmetry for initial adaptation (black) and re-adaptation (green). Shaded patch (gray) indicates split-belt trials. Horizontal line at 0 indicates perfect symmetry. Shaded error bars represent standard error of the mean.

**Supp. Figure 3.**
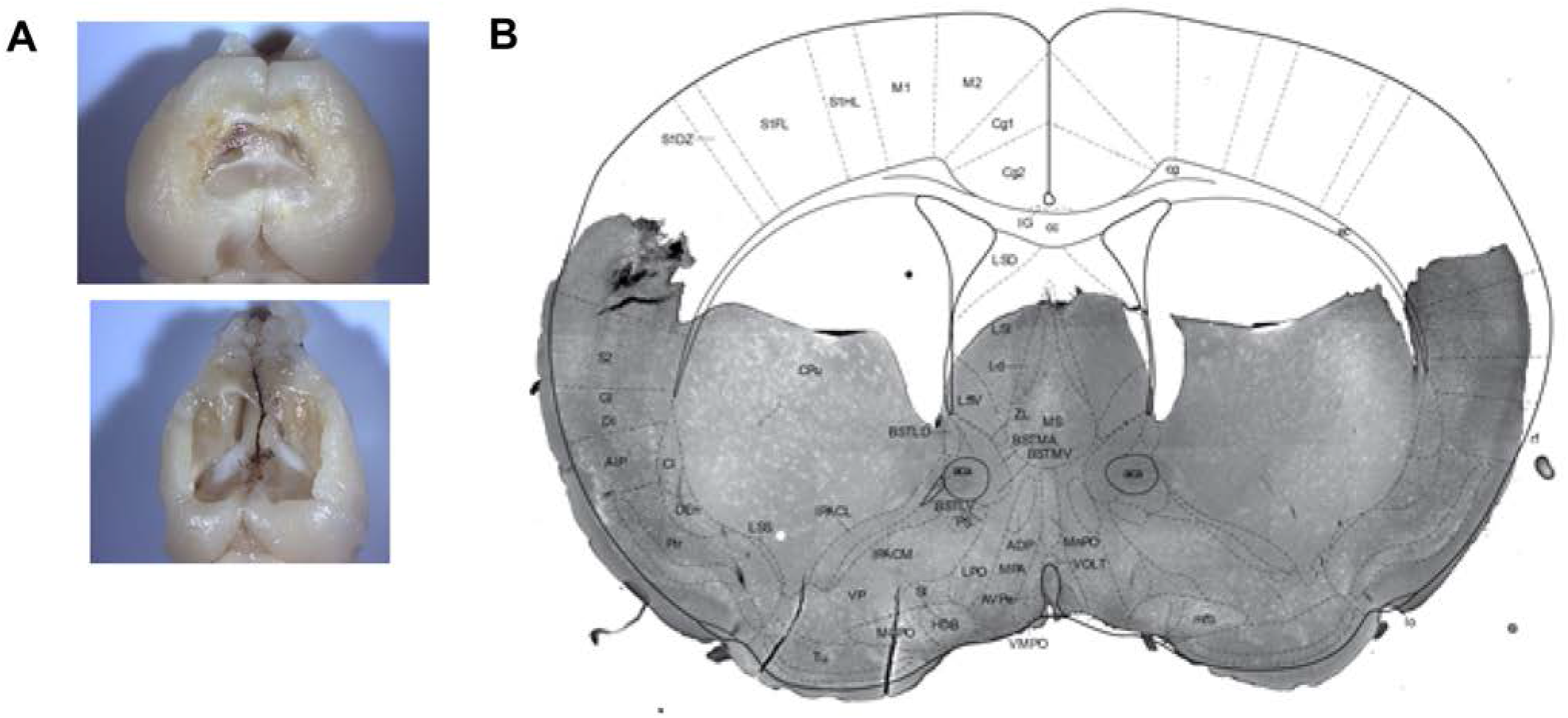
Anatomy of somatomotor cortex lesions. A) Example brains from somatomotor cortex aspiration experiment, showing the extent of aspiration zone for smallest (top) and largest (bottom) lesions. B) Example brain section for somatomotor cortex aspiration experiment overlaid with mouse brain atlas image in coronal view.

**Supp. Figure 4.**
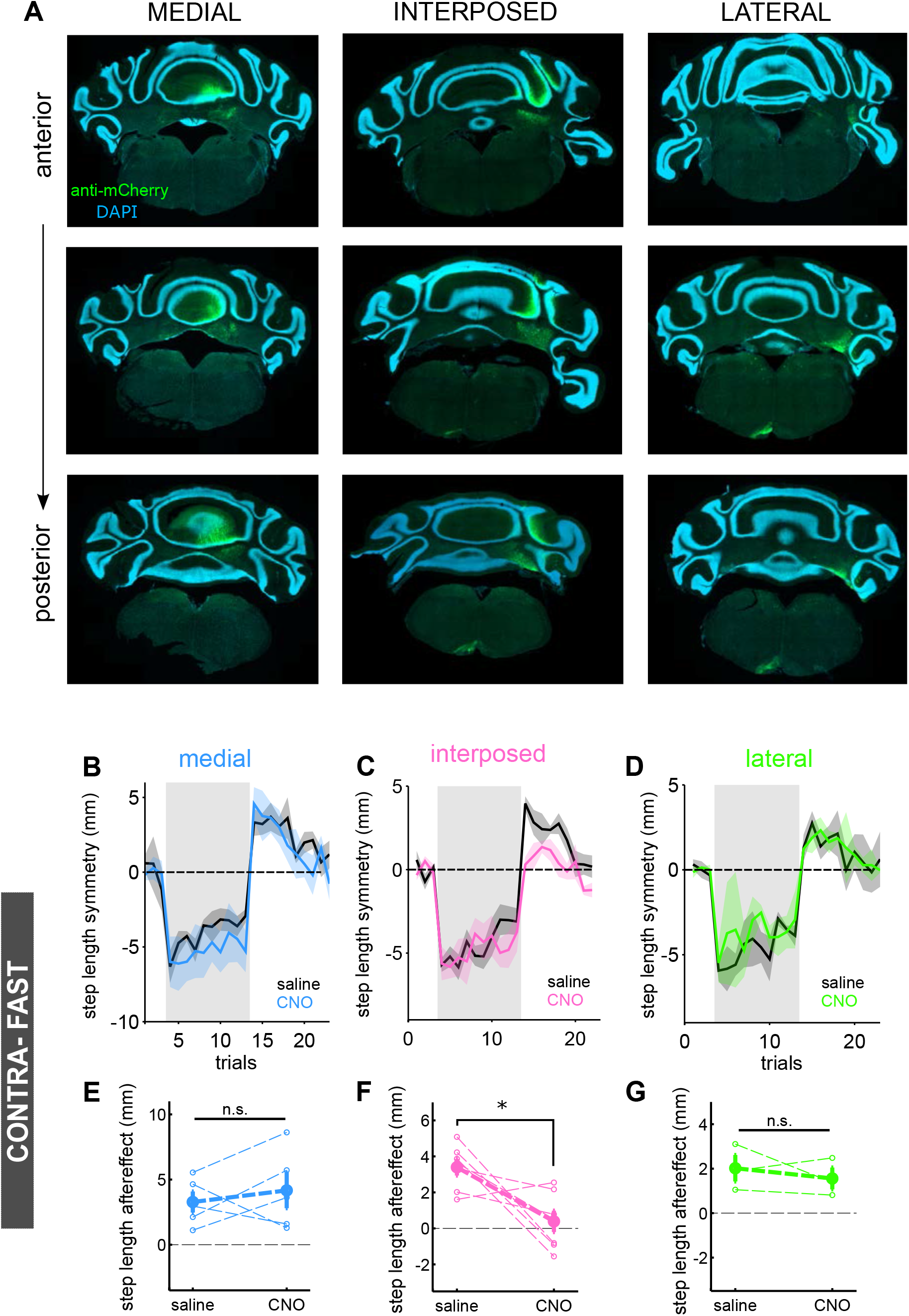
AAV-retro DREADD injections in deep cerebellar nuclei and contra-fast adaptation data. A) Example histology showing anteroposterior spread of virus injections for each of the three deep cerebellar nuclei. (B-D) Average step length adaptation (± s.e.m.) curves for contra-fast medial (B, blue; N = 5), interposed (C,pink; N= 7), and lateral (D, green; N = 3) nucleus injections in with saline (black) and CNO conditions (colors). (E-G) Average step length after-effect sizes (first 2 washout trials) for saline and CNO conditions for medial (E), interposed (F) and lateral (G) nuclei. Individual animals are represented with thin, dashed lines.

**Supp. Figure 5.**
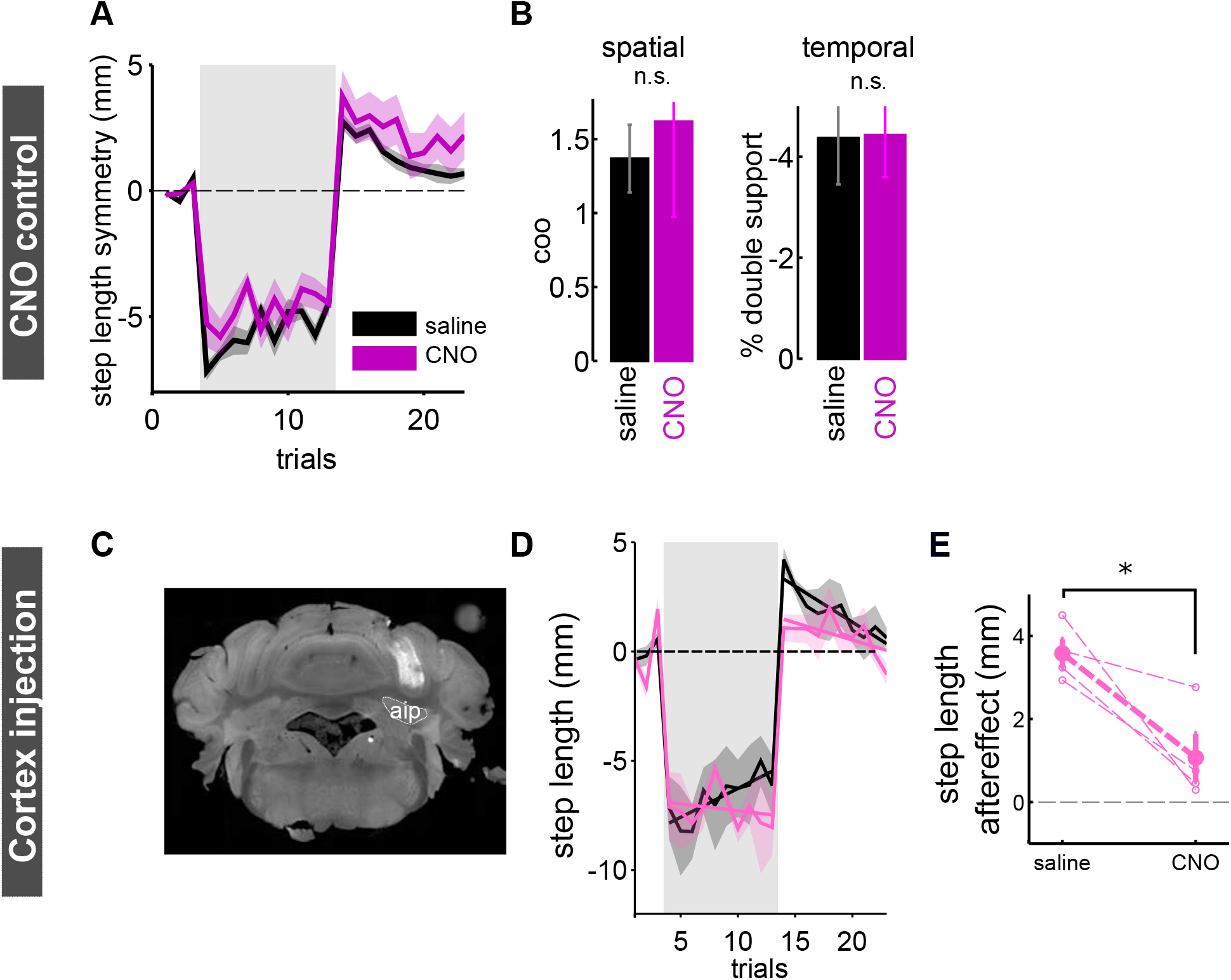
CNO alone does not impair locomotor adaptation and pan-neuronal DREADD injections in intermediate cerebellar cortex are sufficient to impair step-length adaptation. A) Average step length adaptation (± s.e.m.) curves for wildtype animals injected with saline (first adaptation, black) or CNO (second adaptation, magenta). B) Mean spatial and temporal aftereffect (first washout trial) for wildtype mice injected with saline or CNO. Spatial (left) and temporal (right) adaptation were unimpaired by both repeat adaptation and systemic injections of CNO, coo: t_(7)_ =-.51,p = 0.62 ds: t_(7)_= .68, p = 0.52. C) Example cerebellar section from mouse with unilateral injection of pan-neuronal inhibitory DREADD virus in the interposed nucleus. The borders of the interposed nucleus (ip) are indicated by a white, dashed line. D) Average front limb step length symmetry (± SEM) for with saline (black) and with CNO (pink) E) Average step length after-effect sizes (first 2 washout trials) under saline and CNO conditions. Individual animals are shown with thin, dashed lines.

**Supp. Figure 6.**
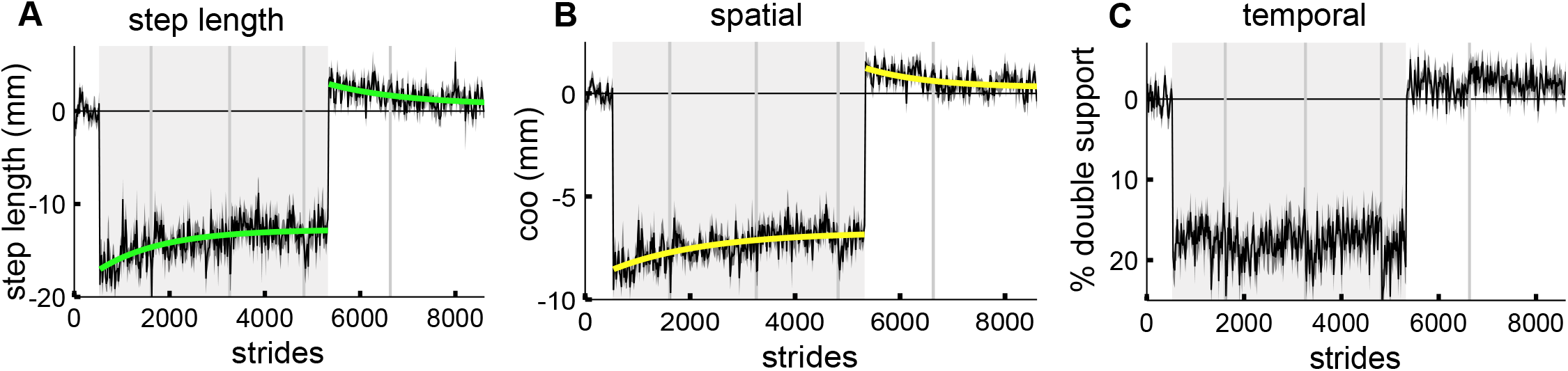
Hind limbs adapt in space but not time. A) Average (n = 17) hind limb step length symmetry (broken down into 10 % trial bins) with exponential decay fits for adaptation and washout periods. Gray shaded patch indicates split-belt trials; darker vertical lines show session breaks B) Same as A for spatial parameter, center of oscillation. C) Same as A for temporal parameter, double support.

**Supp. Movie 1. Example movie of an early split-belt trial with superimposed paw tracks.**

**Supp. Movie 2. Example movie of an interposed injected mouse with and without CNO, walking on tied belts.**

